# Successful asexual lineages of the Irish potato Famine pathogen are triploid

**DOI:** 10.1101/024596

**Authors:** Ying Li, Qian Zhou, Kun Qian, Theo van der Lee, Sanwen Huang

## Abstract

The oomycete *Phytophthora infestans* was the causal agent of the Irish Great Famine and is a recurring threat to global food security^1^. The pathogen can reproduce both sexually and asexually and has a potential to adapt both abiotic and biotic environment^2^. Although in many regions the A1 and A2 mating types coexist, the far majority of isolates belong to few clonal, asexual lineages^3^. As other oomycetes, *P. infestans* is thought to be diploid during the vegetative phase of its life cycle^3^, but it was observed that trisomy correlated with virulence and mating type locus^4^ and that polyploidy can occur in some isolates^5,6^. It remains unknown about the frequency of polyploidy occurrence in nature and the relationship between ploidy level and sexuality. Here we discovered that the sexuality of *P. infestans* isolates correlates with ploidy by comparison of microsatellite fingerprinting, genome-wide polymorphism, DNA quantity, and chromosome numbers. The sexual progeny of *P. infestans* in nature are diploid, whereas the asexual lineages are mostly triploids, including successful clonal lineages US-1 and 13_A2. This study reveals polyploidization as an extra evolutionary risk to this notorious plant destroyer.

---

Understanding the mechanisms of rapid adaption in devastating plant pathogens is of critical importance for disease management. Plant oomycete pathogens employed a flexible reproduction system to gain dual advantages in creating both genotype diversity via sexual crosses and in achieving large population expansion via asexual cycles^2^. Compared to other *Phytophthora* sister species, *P. infestans* has a much larger genome that is enriched in repetitive DNA (∼74%). The genome can be largely divided into gene-dense and gene-sparse compartments, with the latter containing fast-evolving effector repertoire that is required for virulence and host adaptation^7,8^.

An increasing number of reports showed that most *P. infestans* field strains from wide geographical distribution were identified as belonging to a few asexual, clonal lineages^9^. The first recorded clonal lineage HERB-1 was dominant in both Europe and North America between 1845 and 1896, which triggered the Great Irish Potato Famine^5^. Its replacement, the US-1 clonal lineage of the A1 mating type, firstly emerged in North America in the 20th century and dominated the globe^5,10^. Rather recently (1990’s), US-1 was supplanted by more aggressive lineages of the A2 mating type, including 13_A2 in Europe and Asia^11^ as well as US-22 in North America^12,13^. As accumulation of deleterious mutations would lead to rapid extinction of asexual lineages, a phenomenon coined as the Muller’s ratchet^14^, it is puzzling that a few clonal lineages could adapt to a wide geographical area for a considerably long period. What is their common denominator? Although *P. infestans* was known as diploid for decades^3^, several studies indicated trisomy and polyploidy^5,6^. Polyploidy can enhance the vigour in plants and buffer mutational load in asexual reproduction by masking deleterious alleles as reported in both fungi, plants and animals^9,15,16^. It remains unclear to what extent polyploidy contributed to evolutionary advantage of clonal lineages and whether there is link between sexuality and ploidy.

Previously we used multiplex microsatellites to fingerprint 520 *P. infestans* isolates (Supplementary Table 1), including 397 isolates from nine major clonal lineages in four continents and 123 isolates from three sexual populations from the Netherlands, Mexico and Tunisia^9,17,18^. In previous studies, the analysis was only based on the allele size, although differences of allele dosage were observed by peak height. To survey the frequency of polyploidy occurrence and to gain insights into the relationship of reproduction systems and ploidy levels, we re-analyzed the fingerprinting dataset by including scoring of allele peak height, besides molecular size of alleles. The ratio of allele peak heights allows assessing allele dosage at each microsatellite locus (Supplementary Fig.). Surprisingly, we discovered that eight of nine asexual lineages showed high ratio of triallelic loci (0.73-0.99), whereas the value (0.14-0.35) in the three sexual populations is significantly lower (p=0.008). The only exception is the clonal lineage in Northern China (CN-Northern), which is similar to sexual populations in the percentage of triallelic loci. CN-Northern belongs to the A1 mating type and it is a rather old population, as sub-populations evolved in the same lineage^17^. In China, seed potatoes are produced in the North and shipped elsewhere. Strict quarantine is taken for seed tubers to enter the area domestically and abroad. Therefore, the *P. infestans* population there should be rather isolated and remains as diploid.

To further analyze the genome-wide ploidy level, we re-sequenced 11 representative isolates for >50X genome depth from a meta-population from the Netherlands, a major source of international seed potato trade. The meta-population includes three asexual lineages (NL 13_A2, NL 06_A1, and NL 08_A1) and two sexual populations (NL pop 2 and NL pop3, Table 1). Sequences were aligned to the T30-4 reference genome to determine single nucleotide polymorphism (SNP). Relative read depth at each heterozygous SNP was used to determine ploidy level. For diploid genome, the mean of read counts at heterozygous positions should have a single mode at 0.5 (1/2) when the allele ratio is 1:1, while there should be two modes, 0.33 and 0.67 (1/3 and 2/3) when the allele ratio is 1:2 and 2:1 for triploid genomes (Supplementary Table 2). Consistent with the microsatellite fingerprinting analyses, all the nine isolates from the sexual populations displayed diploid mode, whereas the three isolates from the asexual lineages showed triploid mode (Table 2, Fig. 1 and Supplementary Table 2). We also re-analyzed sequences of the seven isolates from a previous report^5^ (Table 2 and Fig. 1), which confirmed that all the four isolates from asexual lineage US-1 and EC-1 showed also triploid mode. These analyses indicate that sexuality and ploidy are strongly associated.

**Table 1.** The allele dosage of microsatellite loci in asexual and sexual populations.

| Lineage / Population | Area | MT<sup>a</sup> | mtDNA<sup>b</sup> | No. isolate | No. locus | 3-allele loci<sup>c</sup> | 2-allele loci<sup>c</sup> | 1-allele loci<sup>c</sup> | 3-allele ratio<sup>d</sup> |
| --- | --- | --- | --- | --- | --- | --- | --- | --- | --- |
| *Asexual lineages* |  |  |  |  |  |  |  |  |  |
| CN Northern | Northern China | A1 | IIa | 67 | 10 | 106 | 213 | 346 | **0.33** |
| CN 13_A2 | Southwest China | A2 | Ia | 64 | 10 | 375 | 25 | 240 | **0.94** |
| CN Fujian | Eastern China | A1 | IIb | 36 | 10 | 153 | 17 | 190 | **0.90** |
| NL 13_A2 | Netherlands | A2 | Ia | 73 | 12 | 591 | 5 | 280 | **0.99** |
| NL 06_A1 | Netherlands | A1 | Ib | 7 | 12 | 52 | 5 | 17 | **0.91** |
| NL 08_A1 | Netherlands | A1 | Ia | 16 | 12 | 91 | 15 | 47 | **0.86** |
| EC-1 | Ecuador | A1 | IIa | 73 | 12 | 422 | 124 | 330 | **0.77** |
| TU-1 | Tunisia | A1 | Ia | 57 | 12 | 312 | 114 | 251 | **0.73** |
| US-1 | US | A1 | Ib | 4 | 12 | 25 | 9 | 14 | **0.74** |
| *Sexual population* |  |  |  |  |  |  |  |  |  |
| NL pop2 | Netherlands | A1, A2 | Ia, IIa | 37 | 10 | 45 | 83 | 241 | **0.35** |
| Mex | Mexico | A1, A2 | / | 44 | 10 | 25 | 154 | 253 | **0.12** |
| TU-2 | Tunisia | A1, A2 | Ia | 42 | 10 | 56 | 105 | 238 | **0.35** |

<sup>a</sup>Mating type
<sup>b</sup>Mitochondria haplotype
<sup>c</sup>3-allele loci means triallelic loci, 2-allele loci means biallelic loci, and 1-allele loci means monoallelic loci
<sup>d</sup>=triallelic loci/(triallelic loci + biallelic loci)

**Table 2.** Ploidy analysis of isolates by genome re-sequencing, flow cytometry and microscope observation.

| Isolate | MT<sup>a</sup> | Lineage / Population | Seq. Depth | DNA Content (pg) | No. chromosome | Ploidy Mode | Reference |
| --- | --- | --- | --- | --- | --- | --- | --- |
| NL00150 | A1 | NL pop3 | 53 | nd<sup>b</sup> | 13.5±1.0 | Diploid | This study |
| NL04092 | A1 | NL pop3 | 54 | nd | nd | Diploid | This study |
| NL04106 | A2 | NL pop3 | 39 | nd | nd | Diploid | This study |
| NL05159 | A1 | NL pop3 | 51 | nd | 14.2±0.8 | Diploid | This study |
| NL05385 | A1 | NL pop3 | 55 | nd | nd | Diploid | This study |
| NL05387 | A1 | NL pop3 | 51 | nd | 13.2±0.9 | Diploid | This study |
| NL05890 | A2 | NL pop3 | 40 | nd | 12.8±0.8 | Diploid | This study |
| NL07434 | A2 | NL pop2 | 54 | 0.56 ±0.02 | 14.2±0.7 | Diploid | Ref. 5 |
| NL08797 | A2 | NL pop2 | 23 | 0.56 ±0.01 | 13.8±0.4 | Diploid | This study |
| P17777 | A1 | nd | 48 | nd | nd | Diploid | Ref. 5 |
| 06_3928A<sup>c</sup> | A2 | NL 13_A2 | 50 | nd | nd | Triploid<sup>c</sup> | Ref. 5 |
| DDR7602 | A1 | US-1 | 13 | nd | nd | Triploid | Ref. 5 |
| LBUS5 | A1 | US-1 | 9 | nd | nd | Triploid | Ref. 5 |
| NL07041 | A2 | NL 13_A2 | 106 | 0.81±0.03 | 20.7±0.4 | Triploid | This study |
| NL08080 | A1 | NL 08_A1 | 43 | 0.82±0.03 | 20.3±0.9 | Triploid | This study |
| NL08452 | A1 | NL 06_A1 | 41 | 0.83±0.02 | 20.8±0.4 | Triploid | This study |
| P13527 | A1 | EC-1 | 20 | nd | nd | Triploid | Ref. 5 |
| P13626 | A1 | EC-1 | 34 | nd | nd | Triploid | Ref. 5 |

<sup>a</sup>Mating type
<sup>b</sup>Not determined
<sup>c</sup>06\_3928A appeared to be tetraploid in previous research<sup>5</sup>. However, our analysis showed it is triploid.

**Figure 1.**
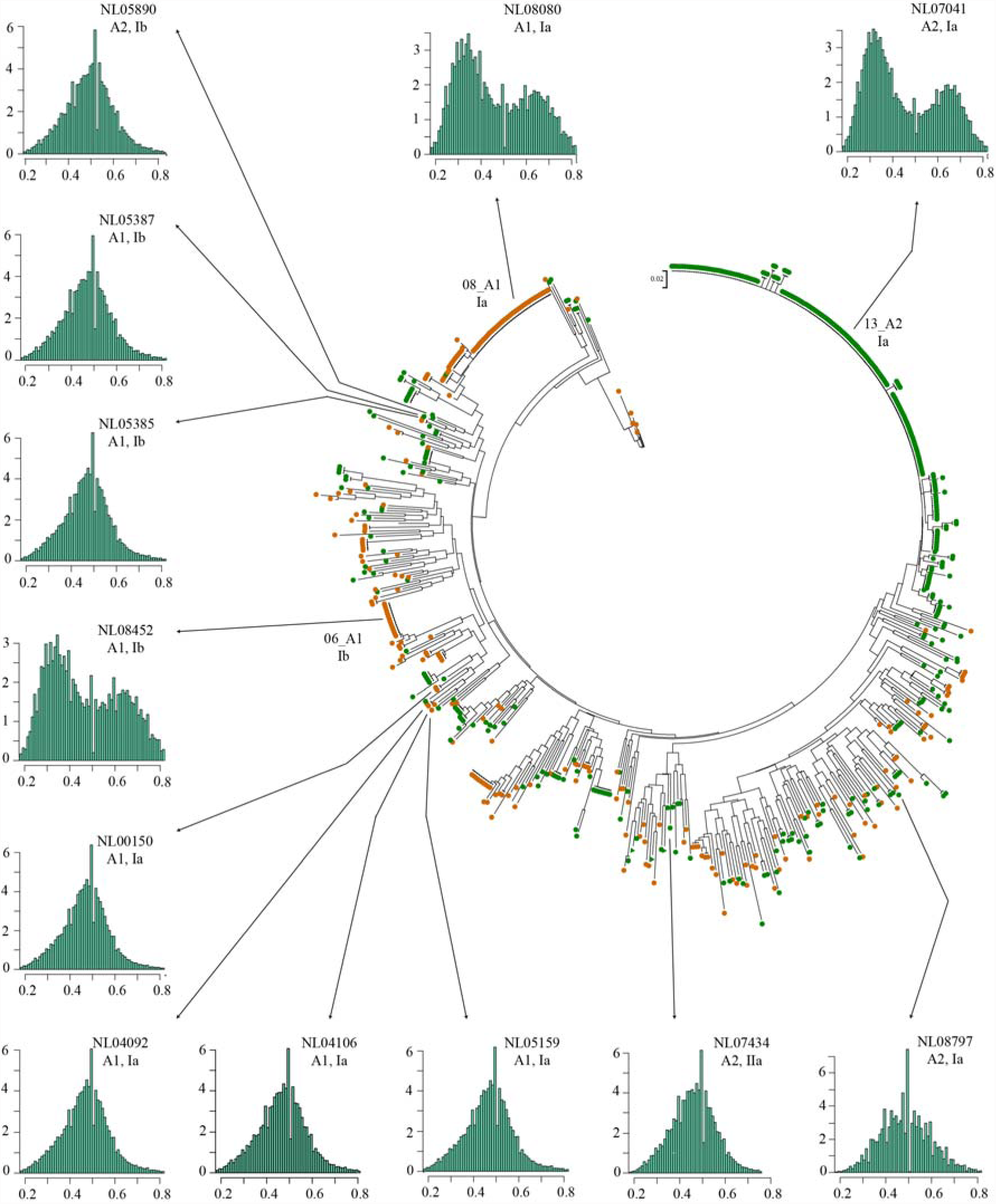
Sexuality and ploidy. The phylogeny tree generated in previous study was used for sampling strategy. Green dots show isolates with A2 mating type, while A1 mating type was marked by orange cycles. Three representative isolates (NL07041, NL08080 and NL08452) were selected from three major clonal lineages in the Netherlands, and nine sexual isolates were from genetic neighbor clades. The ploidy mode was determined with the mean frequency of read counts at heterozygous positions, a single and diploid mode at 0.5, while two modes, 0.33 and 0.67 were for triploid genomes.

In addition to deep re-sequencing, we used flow cytometry and microscope observation to directly evaluate the DNA content and count chromosome number, respectively (Fig. 2). The isolate NL07434 (A2) from the sexual population NL pop2was used as control, since it had been investigated previously^5^ as well as in this study. DNA content of the isolate NL08797 (A2) from NL pop 2 is the same as NL07434. As expected, DNA content of the three asexual isolate NL08080 (A1), NL08452 (A1), and NL07041 (A2) is about 1.5 times of that of NL07434 and NL08797, confirming that they are triploid. The average of chromosome number for isolates from sexual populations is 13.6, whereas 20.6 for isolates from asexual lineages (Table 2, Supplementary Table 3). DNA content measurement and chromosome counting further support that these asexual isolates are triploid.

**Figure 2.**
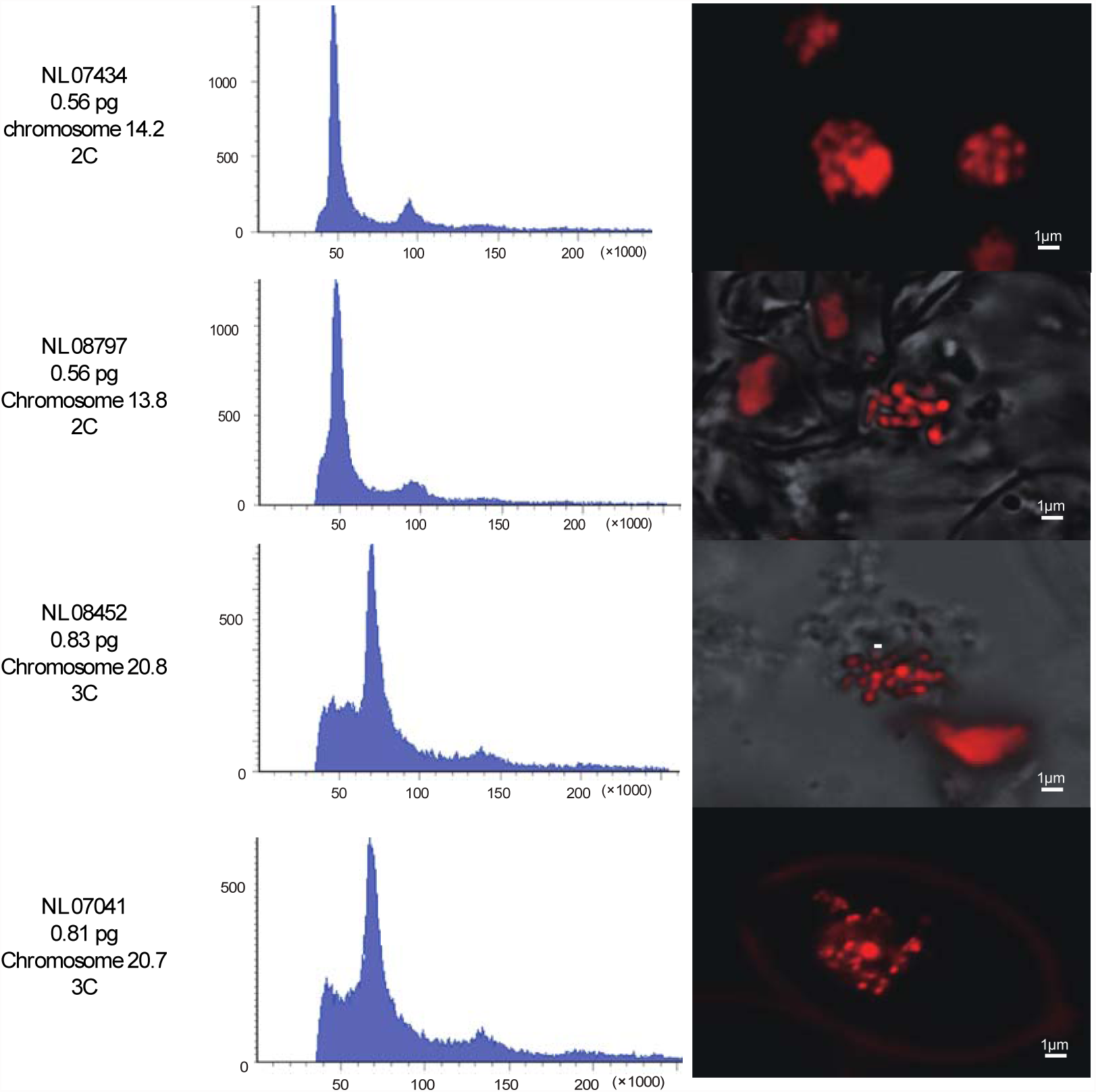
DNA content and chromosome number. Two isolates from asexual lineage (NL08452 and NL07041) and two isolates from sexual populations (NL07434 and NL08797) were investigated. DNA content was determined by flow cytometry. The chromosome number was counted by adjusting the focal levels on Laser Scanning Confocal Microscope (LSM780, Zeiss).

In summary, *P. infestans* can no longer be considered to be only diploid, which should be taken into account in future studies. The observed triploidy must play an imperative role in the successful global epidemic of modern asexual lineages such as US-1, NL 13_A2, TU-1, NL 06_A1 and NL 08_A1. Polyploidization in *P. infestans* is likely to enhance fitness, as reported in yeast that tetraploid had a higher beneficial mutation rate than haploid and diploid strains^19^. Ancient or old asexual lineages such as HERB1 and CN Northern are diploids, therefore it is worth further investigation on when and how asexual lineages became triploid. Was the transition of the ploidy level due to survival pressure from disease resistance genes that were incorporated into potato breeding after the Great Famine or linked to the frequent use of pesticide deployed in the last 50 years to control this disease? This study revealed a new dimension of genomic feature of the great evolutionary risk of *P. infestans*, which should also be considered in future agricultural management.

## Methods

### Definition of asexual and sexual populations

This study only focuses on field isolates. We adopted the definition of clonality widely accepted in papers dealing with the population structure of pathogens^20^. The definition of asexuality (clonal lineage) and sexuality (sexual population) in this study do not refer to the cytological mechanism of reproduction, but rather to the population structure that results from an absence or restriction of genetic recombination^20^. The asexuality obtains wherever isolates with the same mating type show multilocus genotypes (MLGs) that are identical or nearly identical. In contrast sexual progeny shows extensive recombination of alleles.

### Isolate collection

In this study, isolates from China^17^, The Netherlands^9^, Ecuador^18^, Tunisia (manuscript prepared), and Mexico were investigated (Table 2). The clonal lineages and sexual populations have been defined by MLGs in those previous studies. The clonal lineages and sexual populations used in this study have been defined by MLGs in the previous studies.

### Microsatellite analysis

Twelve microsatellite markers were used^21^-^24^. Amplification of the SSR markers was carried out as described by Li et. al^24^. The amplicon was capillary electrophoresed on an automated ABI 3730 according to the manufacturer's instructions. SSR allele sizing was performed using GeneMapper v3.7 software (Applied Biosystems, USA).

### Re-sequencing data and SNP calling

The reference genome sequence of the artificial strain T30-4 and the re-sequencing data used in this study were published in previous studies^5,7,10,11^. The method of read mapping was used as described^5^. The re-sequenced isolates selected from previous studies were analyzed again here. Since the reference genome has 4921 supercontigs, we only analyzed the first 100 longest supercontigs.

The isolates were sequenced using the Illumina Hi-Seq 2000 sequencer. The sequencing averagely generated 100 million 100-bp paired-end reads for each isolate. The sequencing reads of each isolate were mapped to the T30-4 reference genome using BWA^25^ and SNP calling was conducted subsequently using Sequence Alignment/Maptools^26^. Several criteria was considered in SNP filtering: (1) a SNP should be bi-allelic between the isolate and T30-4 genomes; (2) at the SNP locus, the phred quality score of base sequencing and score of read mapping should be both higher than 30; (3) each allele of a SNP should be supported by at least 4 reads; (4) the non-reference allele frequency should be between 0.2-0.8 at a heterozygous SNP locus.

### Flow cytometry

The nuclei were collected by simply chopping hyphae with a scalpel blade in cold phosphate buffered saline (PBS)^27^. Another method used was zoospore cultures. Zoospores were induced at 4⍰ for two hours and harvested in excess of 10^4^ spores ml^-1^. The nuclei were stained by 10ug/ml propidium iodide (Sigma). The samples were delivered into a laser BD FACSCalibur Flow Cytometer for data analysis.

### Microscopy observation of chromosomes

The young hyphae stained by 50ug/ml propidium iodide (Sigma) were transferred to a microscope slide for observation. The observation was preformed with Zeiss LSM 780 Laser Scanning Confocal Microscope.

## Supplementary Information

includes one figure and three tables and is available in attached files.

## Acknowledgements

We thank G. J. T. Kessel and G. B. M. van den Bosch (Wageningen University) for providing *P. infestans* isolates. We appreciate Z. Zhang, B. Xie and S. Dong for useful discussion and E.Jacobsen (Wageningen University) and T.Städler (ETH Zurich) for critical reading. This work was supported by Chinese Special Fundfor Agro-scientific Research in the Public Interest (201303018), National High Technology Research and Development Program (2013AA102603), National Natural Science Foundation of China (31201257), Ministry of Finance in China (1251610601001), Science and Technology Innovation Program of Chinese Academy of Agricultural Sciences (CAAS-ASTIP-IVFCAAS).

## Author Contributions

Y. L. and S. H. designed the study and wrote the paper. Y. L. performed SSR analysis and chromosome observation. Q. Z. analysed re-sequencing data. Q.K. performed flow cytometry. T. L. provided the raw SSR data.

## Author Information

The authors declare no competing financial interests. Correspondence and requests for materials should be addressed to Y. L. and S. H.

